# EGLN Inhibition Reduces Gastrointestinal Radiation Toxicity and Improves Survival in a Murine Model of Locally Advanced Pancreatic Cancer

**DOI:** 10.1101/195610

**Authors:** Tara N. Fujimoto, Lauren E. Colbert, Jessica M. Molkentine, Laura Baseler, Amit Deorukukhar, Charles V. Kingsley, Ramesh C. Tailor, Gabriel O. Sawakuchi, Cullen M. Taniguchi

## Abstract

Locally advanced pancreatic cancer (LAPC) almost always fatal since it is unresectable and chemotherapy is only modestly effective. The efficacy of radiation therapy (RT) for LAPC is limited by the potentially fatal toxicity to nearby intestines. There are no FDA-approved medications that can prevent this radiotoxicity, but we find that FG-4592, a small molecule inhibitor of EGLN proteins, significantly reduces radiation damage to the intestines without radioprotecting tumors. KPC (KrasLSL/+; Trp53FL/+; Ptf1aCre/+) animals received dose-escalated radiation treatments with and without FG-4592 for radioprotection. High-dose RT reduced death from local progression, improved survival, and shifted the patterns of failure to a late metastatic death compared to controls. The addition of FG-4592 to RT further improved survival compared to vehicle controls by eliminating radiation-induced gastrointestinal toxicity. Thus, selective protection of the intestinal tract by EGLN inhibition may enable higher, and potentially definitive doses of cytotoxic therapy to be delivered to LAPC.

**One Sentence Summary:** The EGLN inhibitor FG-4592 allows higher, and potentially definitive, doses of radiation to be delivered to pancreatic cancer by reducing normal tissue toxicity without protecting tumors.

## Introduction

Pancreatic cancer portends a poor prognosis, and despite ever increasing knowledge of its biology (*1*), this disease is projected to the be the second leading cause of cancer-related death by 2030 (*2*). Definitive treatment for pancreatic cancer is extremely important since the primary tumor can kill the patient from enxtension and obstruction of nearby bile ducts, blood vessels and visceral organs even before the disease has a chance to metastasize (*3*). This is a particular problem for patients with locally advanced pancreatic cancer (LAPC), since their tumor is by definition not surgically resectable, but not yet metastatic. Thus, for patients with LAPC, radiation therapy is often the only available treatment option for local control.

Radiation therapy can be efficacious against LAPC, but only if the tumor receives a radiobiological equivalent of 70Gy or more (*4*), which is almost never achieved because most pancreatic tumors grow too close to the duodenum, jejunum, or stomach, which are inherently radiosensitive organs that can only tolerate an equivalent of 50Gy. Doses that exceed this limit can cause morbid complications such as intestinal bleeding, visceral perforation, or fistulas (*5, 6*). Recent attempts to improve radiation therapy with conformal approaches such as intensity-modulated radiation therapy (IMRT) or heavy ion therapy (*7*) have improved toxicity (*8*), but are still unable to deliver a definitive dose due to the fact that the duodenum and jejunum often cannot be shielded or protected sufficiently from the high-dose treatment penumbra due to its anatomical proximity to the pancreas. Thus, treatment-related gastrointestinal (GI) toxicity may be the single greatest barrier to the delivery of definitive treatments for unresectable pancreatic cancer. Unfortunately, there are no known medications that can selectively protect the intestines from radiation toxicity.

The EGLN family of α-ketoglutarate-dependent prolyl hydroxylases controls the cellular response to hypoxia by directing the destruction of the hypoxia-inducible factor (HIF) family of transcriptions factors under normal oxygen tension (*9*). When the EGLN proteins are inhibited genetically or pharmacologically, HIF can be stabilized in normoxic tissues, which allows the potentially therapeutic benefits of this pathway to exploited. For instance, we and others have previously demonstrated that the genetic or pharmacologic pan-inhibition of the three mammalian isoforms of EGLN (EGLN1-3) mitigated and protected animals from otherwise lethal doses of gastrointestinal radiation (*10, 11*). Although these phenotypes were robust, these studies were not yet suitable for clinical translation because the radiation fields were toxic by design (*12, 13*) and the EGLN inhibitor used did not have pharmacokinetics suitable for human use (*14*). It was unclear whether EGLN inhibitors could be used along with fractionated radiation for pancreatic cancer since the large radiation fields and doses were too toxic for direct translation to the clinic.

Here, we demonstrate that FG-4592, a clinically and pharmacologically relevant pan-EGLN inhibitor (*15*), significantly reduces gastrointestinal toxicity from high-dose fractionated radiation treatments. This radioprotection did not affect tumor growth and actually increased lifespan of treated mice by reducing both tumor-related and treatment-related deaths, which provides a proof of concept for this approach in patients.

## Results

### The EGLN inhibitor FG-4592 Significantly Reduces Gastrointestinal Toxicity from Fractionated Radiation

Current preclinical models that use a single fraction of whole body or whole-abdominal radiation would never be used clinically due to a subtherapeutic dose (14-20Gyx1,(*16*)) and high levels of radiation-induced toxicity (*12, 13*). In order to find a regimen that better resembled an effective dose in clinical studies (*4*), we tested the tolerability of a limited subdiaphragmatic 15mm upper abdominal field (UA-XRT, Fig 1A) guided by daily cone beam CT imaging sufficient for isocenter localization. This field size was chosen since it would be large enough to treat most pancreatic tumors, while not excessively radiating other abdominal organs such as the kidneys. Of note, this field does treat a portion of the liver, stomach and jejunum along with the entire pancreas and duodenum.

**Fig 1.**
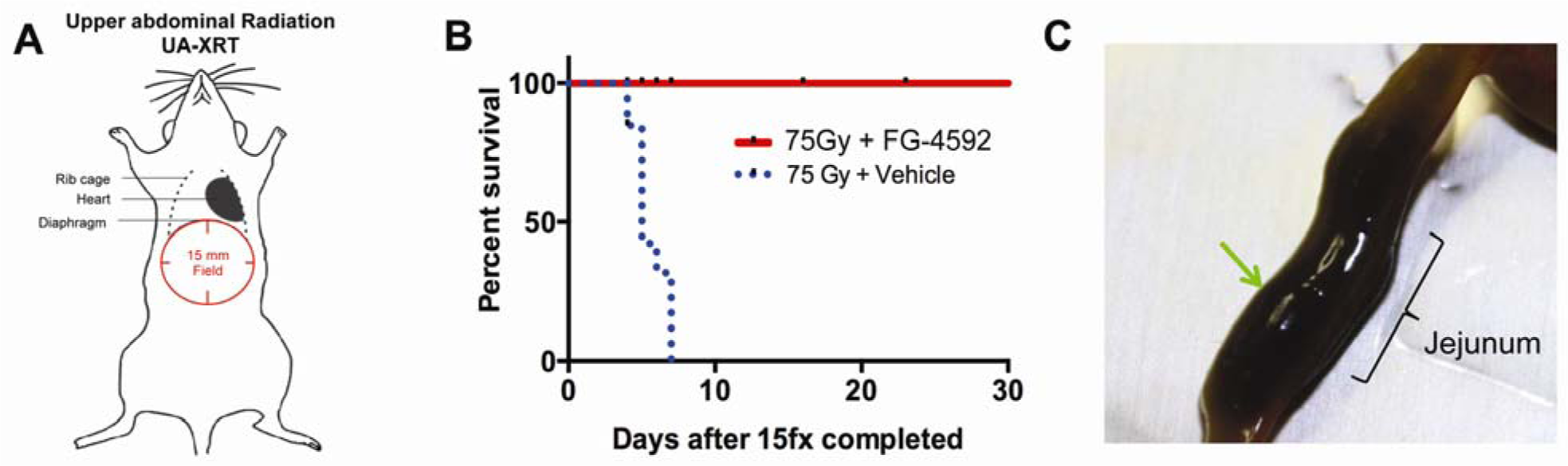
(**A**) Upper abdominal radiation (UA-XRT) (**B**) Improved survival of mice treated with FG-4592 after 75Gy/15fx of UA-XRT (n=10/tx cohort) (**C**) Example of digested blood within jejunum (green arrow).

To determine whether EGLN inhibition could protect mice from this high-dose, clinicially-relevant form of fractionated radiation, we subjected wild-type, non tumor-bearing C57BL/6J mice to 75Gy in 15 fractions (BED_10_=112.5Gy) directed to an UA-XRT field with FG-4592 or vehicle control for radioprotection (see Methods for full details). This 15-fraction regimen is similar to the dose and fractionation used at UT MD Anderson Cancer (*4, 8*) and other institutions (*17*) for locally advanced pancreatic cancer. Strikingly, 100% of the mice that received the cytoprotectant FG-4592 were alive 30 days after treatment, while none of the vehicle mice lived beyond 10 days (Fig 1B). Necropsies of the vehicle controls revealed guaiac positive stools, suggesting a bleeding event from radiation, which is consistent with clinical toxicities (Fig 1C). There was no evidence of bleeding in mice treated with FG-4592.

### FG-4592 Selectively Enhances HIF Expression in Intestines, but not Tumors

A radiation protector should have a therapeutic window that preferentially affects normal tissues, but not tumors. Thus, we treated C57BL/6 mice with a single oral dose of FG-4592 and found a ten-fold increase in the expression of both HIF1 and HIF2 in the duodenum and jejunum (Fig 2A). To understand whether FG-4592 would also stabilize HIF levels in tumors, we bred mice that harbored a conditional activated *<u>K</u>ras* allele (G12D) and a heterozygous *Tr<u>p</u>53* loss-of-function allele driven by a pancreatic specific <u>C</u>re (*Ptf1a*-Cre) (*18*), also known as KPC mice. A small cohort of KPC mice with spontaneous tumors were treated with FG-4592 or 75Gy of RT in 15 fractions with or without FG-4592 to determine how these treatment might alter HIF expression in tumors. There was significant expression of HIF1 and HIF2 at baseline in untreated tumors and the addition of either RT or FG-4592 did not further increase this expression of either HIF isoform (Fig 2B).

**Fig 2.**
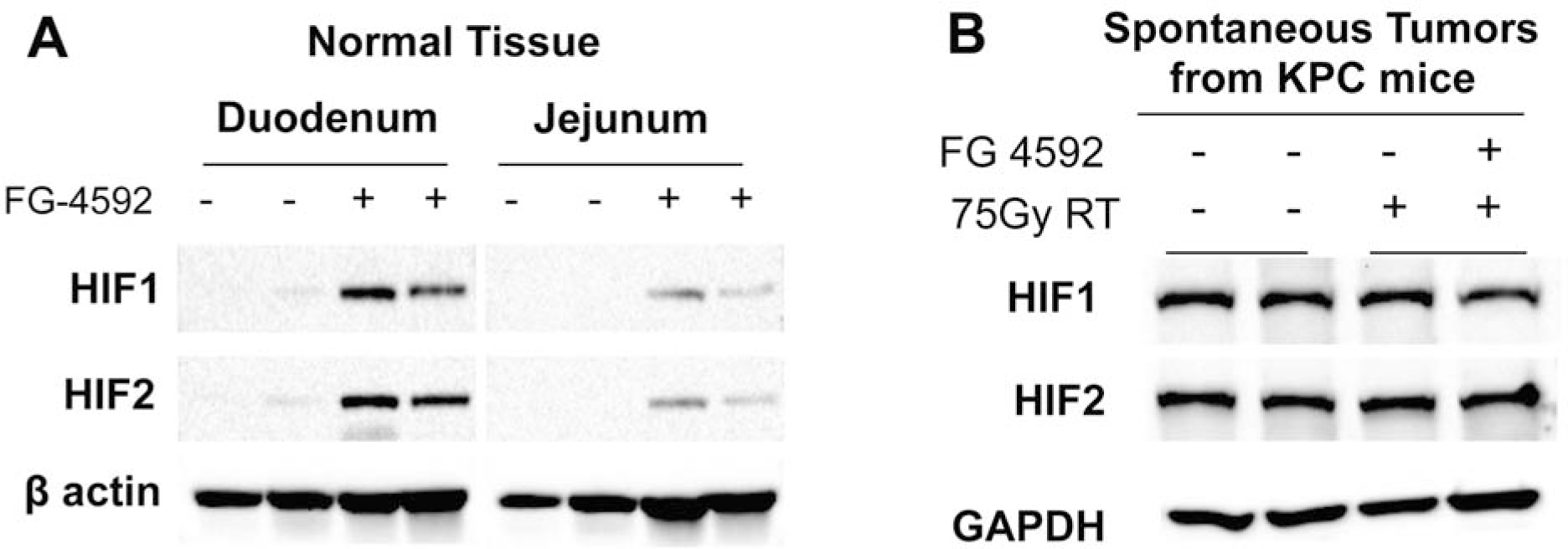
FG-4592 Enhances HIF expression in the intestine, but not in tumors. (A) Western blots demonstrating HIF expression in the duodenum and jejunum 6 hours after receiving FG-4592. (B) EGLN inhibition by FG-4592 nor RT alters HIF expression within tumors

### Radiation Therapy with EGLN Inhibition Improves Survival in KPC Mice

To test whether the addition of definitive radiation therapy with or without the radioprotector FG-4592 could improve outcomes in pancreatic cancer, we initiated a preclinical trial using KPC mice with a hetereozygous deletion of *Trp53* (*Kras*^LSL/+^; *Trp53*^FL/+^; *Ptf1a*^Cre/+^). These animals were chosen because previous reports had demonstrated that this model presented with locally advanced disease (*19*) instead of widespread metastases, which common in models that use the *Trp53^R172H^* mutant knock-in construct (*20*). Moreover, our KPC animals were backcrossed to C57BL/6 genetic background over 10 generations to reduce potential variability in normal tissue radiation responses that often occurs in mixed backgrounds (*21*). The experimental schema is shown in Fig 3A. KPC mice with the appropriate genotype were screened for tumors by weekly ultrasounds beginning at 16 weeks of age. When a tumor was identified, mice were sequentially assigned to receive one of four possible treatments: Vehicle only (VEH), FG-4592 (FG), RT concurrent with vehicle (RT+VEH), or RT concurrent with FG-4592 (RT+FG). Mice were enrolled on a rolling basis and the total time from diagnosis to treatment end was approximately three weeks. Radiation treatments were planned to administer a total of 75Gy in 15 daily fractions, which is a dose that has been correlated clinically with better outcomes. Mice who received >65Gy were considered as receiving full dose RT and completing treatment. During treatment, mice were monitored twice daily and any animals that met criteria for euthanasia were sacrificed and subjected to a blinded necropsy under the supervision of a veterinary pathologist, who also assigned causes of death.

**Fig 3.**
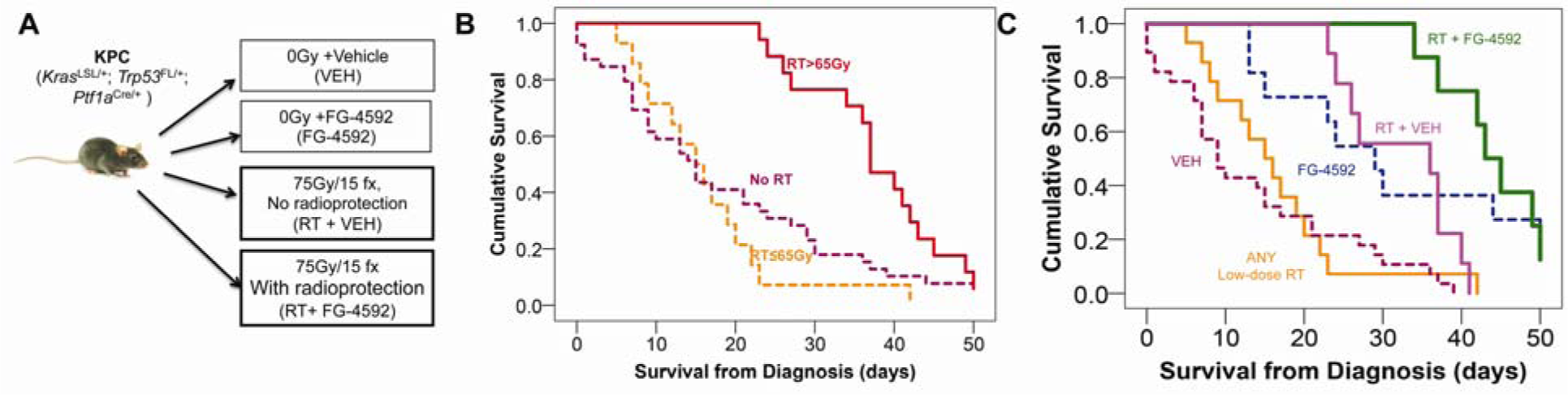
High-Dose Radiation with Radioprotection Improves Survival. **(A)** Experimental scheme for KPC animals. Mice were enrolled to receive no RT (0Gy) with either vehicle or FG-4592 or to dose-esclated radiation (planned 75Gy/15fractions UA-XRT) with vehicle or FG-4592 for radioprotection. (**B**) Kaplan-Meier survival curves for groups divided by radiation treatment dose. Animals receiving RT>65Gy demonstrated the highest overall survival (red solid line, P=0.005), compared to ≤65Gy RT (orange dashed lines) or no RT (Vehicle or FG-4592 only (purple dashed line) **(C)** Kaplan-Meier survival curves for all animals, with highest overall survival for mice treated with RT + FG-4592 (p<.0001)

Seventy mice were evaluated and their characteristics are described in Table 1. A total of 40 mice were assigned to an arm without radiation, which included vehicle treatment alone (N=28) or oral FG-4592 alone (N=11). Due to disease progression, 10% (4/41) of animals in the VEH group and 34% (10/29) of the animals in the FG group were not able to complete the full course of treatment. Thirty mice were assigned to high-dose fractionated radiation, with vehicle (N=13) or concurrent FG-4592 given for radioprotection (N=18). Despite the higher planned doses of radiation, only 69% (9/13) of the animals from the RT+VEH group and 44% (8/18) animals from RT+FG group could complete the treatments, due to progression of disease during treatment.

**Table 1.** Distribution of Tumor and Treatment Characteristics for All Enrolled Animals by Treatment Group

| **RADIOPROTECTANT** | **VEHICLE (N=41)** |  |  | **FG-4592 (N=29)** |  |  |
| --- | --- | --- | --- | --- | --- | --- |
| **RT DOSE** | **No RT<sup>1</sup>**<br>**(N=28)** | **<=65Gy<sup>1</sup>**<br>**(N=4)** | **>65Gy<sup>1</sup>**<br>**(N=9)** | **No RT<sup>1</sup>**<br>**(N=11)** | **<=65Gy<sup>1</sup>**<br>**(N=10)** | **>65Gy<sup>1</sup>**<br>**(N=8)** |
| **SEX** |  |  |  |  |  |  |
| **FEMALE** | 10 (36) | 3 (75) | 8 (89) | 8 (73) | 3 (30) | 2 (33) |
| **MALE** | 18 (64) | 1 (25) | 1 (11) | 3 (27) | 7 (70) | 6 (67) |
| **TUMOR LOCATION** |  |  |  |  |  |  |
| **BODY** | 10 (36) | 0 (0) | 5 (56) | 5 (45) | 4 (40) | 5 (63) |
| **HEAD** | 13 (46) | 1 (25) | 3 (33) | 6 (55) | 3 (30) | 1 (13) |
| **TAIL** | 5 (18) | 3 (75) | 1 (11) | 0 (0) | 3 (3) | 2 (25) |
| **CAUSE OF DEATH** |  |  |  |  |  |  |
| **LOCAL TUMOR** | 21 (75) | 4 (100) | 1 (11) | 6 (55) | 6 (60) | 3 (38) |
| **METASTATIC** | 5 (18) | 0 (0) | 3 (33) | 5 (45) | 4 (40) | 5 (62) |
| **RADIATION TOXICITY** | 0 (0) | 0 (0) | 5 (55) | 0 (0) | 0 (0) | 0 (0) |
| **MISSING** | 2 (7) | 0 (0) | 0 (0) | 0 (0) | 0 (0) | 0 (0) |
| **JAUNDICE** |  |  |  |  |  |  |
| **NO** | 18 (64) | 4 (100) | 8 (89) | 7 (64) | 10 (100) | 7 (88) |
| **YES** | 8 (29) | 0 (0) | 0 (0) | 2 (18) | 0 (0) | 0 (0) |
| **MISSING** | 2 (7) | 0 (0) | 1 (11) | 2 (18) | 0 (0) | 1 (13) |
| **AGE AT DIAGNOSIS**<br>**(DAYS)<sup>2</sup>** | 21 (12-29) | 26 (21-32) | 23 (12-27) | 18 (14-28) | 19 (13-36) | 27 (19-32) |
| **WEIGHT AT DIAGNOSIS**<br>**(GRAMS)<sup>2</sup>** | 21 (12-29) | 24 (24-25) | 23 (20-26) | 23 (20-28) | 25 (20-30) | 28 (26-28) |
| **TUMOR VOLUME AT**<br>**DIAGNOSIS (MM<sup>3</sup>)<sup>2</sup>** | 101 (10-371) | 213 (51-371) | 63 (9-219) | 66 (25-109) | 91 (20-288) | 282 (114-587) |
| **TUMOR VOLUME AT**<br>**DEATH (ULTRASOUND -**<br>**MM<sup>3</sup>)<sup>2</sup>** | 367 (19-3958) | 735 (51-1222) | 288 (23-1620) | 380 (103-1004) | 277 (97-605) | 282 (114-587) |
| **WEIGHT AT DEATH**<br>**(GRAMS)<sup>2</sup>** | 27 (25-28) | 24 (23-24) | 19 (16-20) | 21 (17-28) | 22 (16-26) | 19 (15-27) |

The administration of high-dose radiation therapy of >65Gy (high-dose) with or without radioprotection improved survival compared to all animals that received ≤65Gy (low-dose) of treatment or no RT (Fig 3B; 37 days [95% CI 33-41] vs. 15 days [10-21] vs. 15 days [10-20], respectively; p=0.005). The median overall survival was compared amongst the treatment groups and was highest (Fig 3C; p<.001) for mice that received RT with FG-4592 for radioprotection (43 days, [95% CI 39-47]) versus RT with vehicle (36 days, [95% CI 10-62]), FG alone (29 days, [95% CI 21-37]), or vehicle alone (9 days, [95% CI 5-13]).

### High-Dose Fractionated Radiation Improves Survival and Reduces Symptoms of Local Progression

Study mice were monitored twice daily and subjected to euthanasia with necropsy analysis when predefined criteria for disease burden or symptoms were met (see Methods). Each necropsy was directly supervised by a veterinary pathologist, who analyzed the tissues and carcasses and assigned a probable cause of death while blinded to treatment. From these necropsies, we were able to understand how the patterns of spread changed with and without radiation therapy. Death from local progression were usually instantly apparent upon necropsy, usually in the form of gastric or intestinal obstruction as depicted in Fig 4A. Even if metastases were present, they were not held responsible for death if there was an otherwise obvious obstruction. Death from progression of the primary pancreatic tumor occurred in 69% (27/39) of animals that did not receive radiation therapy (vehicle alone or FG-4592 alone) compared to 17% (3/17) of animals that received full dose pancreas-directed radiation therapy (Fig 4B; p<.0001). Locally advanced pancreatic tumors also frequently causes obstructive jaundice, which is a highly morbid condition that can lead to biliary sepsis and death (REF). This symptom was frequently seen in advanced disease in our KPC animals, as illustrated in Fig 4C. Obstructive jaundice was present in 64% (25/39) of mice who did not receive RT versus 0% (0/29) who received any dose of RT with or without radioprotection (Fig 4D; p=0.02).

**Fig. 4.**
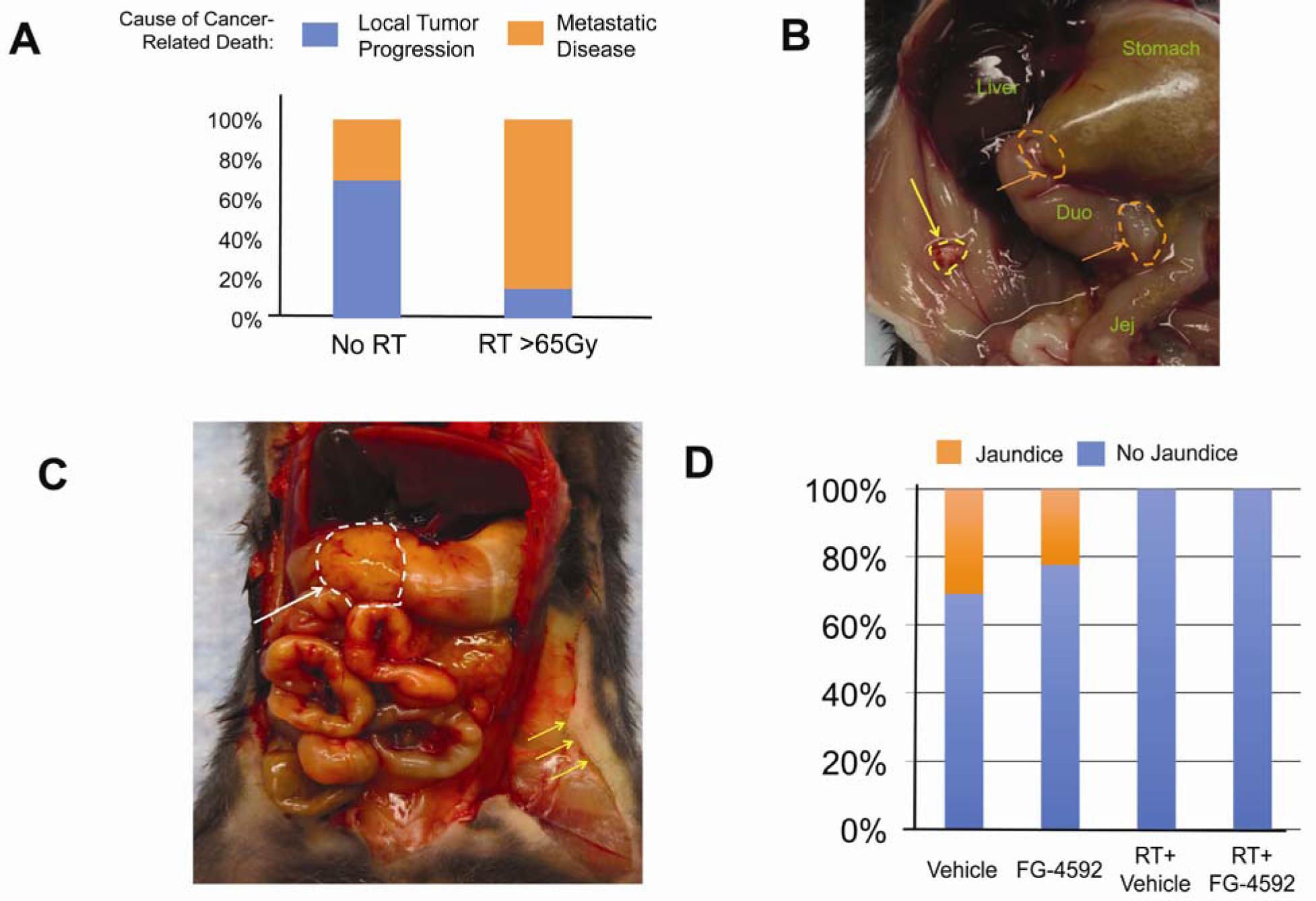
High-dose RT reduces symptoms caused local progressionof pancreatic cancer. (**A**) Cause of death analysis demonstrating lower death due to local tumor burden in >65Gy radiation treatment groups (vehicle or FG-4592) vs no RT treatment groups (P<.001) (**B**) Necropsy photos showing acute and fatal obstruction at the gastric antrum and duodenum (orange dashed lines). Of note, despite there being a peritoneal tumor deposit, the cause of death was local progression causing obstruction. **(C)** Locally obstructive pancreatic tumor (white dahes lines and arrow) causing obstructive jaundice (yellow arrows). **(D)** Higher rates of jaundice in animals who did not receive local radiation therapy (p=0.02)

### EGLN Inhibition Ameliorates Radiation-related Mobidity and Mortality

High-dose radiation with radioprotection provided by FG-4592 provided the greatest survival benefit of all the study cohorts. To understand why, we looked at all causes of mortality, including deaths caused by any treatment. As predicted from prior studies in animals with our KPC genotype (*Kras*^LSL/+^; *Trp53*^FL/+^; *Ptf1a*^Cre/+^), animals that received no direct treatment to their primary tumor died chiefly from local progression (*19*). Local progression was the chief cause of death in 75% (21/28) of KPC animals that received vehicle and 55% (11/16) of mice that received FG-4592 alone. In contradistinction, 55.6% (5/9, Fig 5C) of animals that received high-dose RT to the upper abdomen without radioprotection died from fatal GI bleeding, whereas no bleeding was seen in animals who received high dose RT+ FG-4592 (Fig 5D; P<.0001). An example of such bleeding caused by radiation toxicity is shown in Fig 5E.

**Fig. 5.**
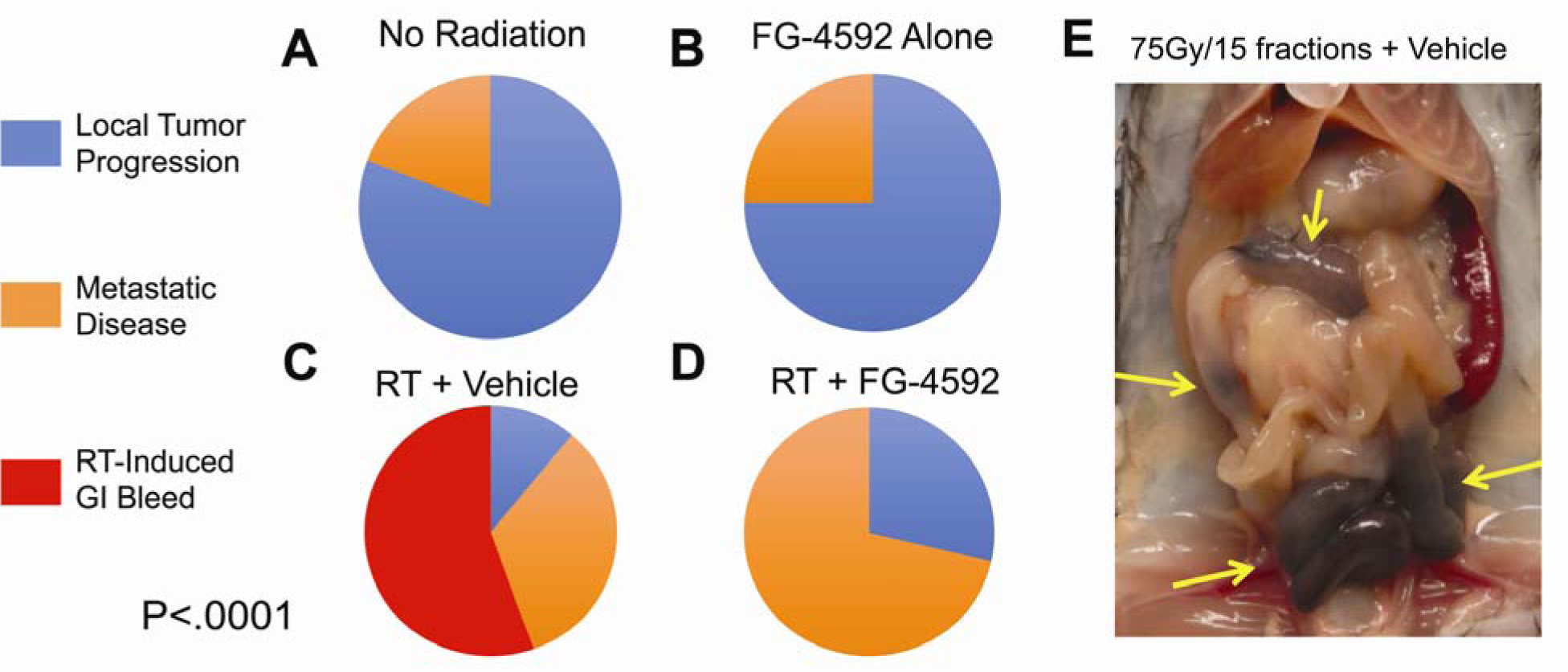
Radioprotection with FG-4592 reduces radiation-induced gastrointestinal toxicity. Distribution of all causes of death from necropsy for (A) Vehicle alone, (B) FG-4592 alone, (C) >65Gy RT + Vehicle and (D) >65Gy RT + FG-4592, with an example of gastrointestinal bleeding noted on necropsy from a mouse that received 75Gy of fractionated radiation without radioprotection (yellow arrows, (E)).

## Discussion

High-dose radiation therapy improved survival and reduced morbidity from pancreatic cancer in this preclinical model by a mechanism that is illustrated in Fig 6. When no locally directed therapy is given, mice die rapidly from a locally destructive primary tumor (Fig 6A), as has been previously reported in this model (*19*). Treatment of the primary tumor with high-dose radiation therapy reduces clinical symptoms of local tumor progression such as jaundice (Fig 4C) and improves survival (Fig 3B). These reductions in mortality were counterbalanced by a high incidence of treatment-related toxicity in those mice that were not radioprotected (Fig 5C, and 6B). FG-4592, on the other hand, ameliorated radiation enteritis and maximized survival (Fig 3C and 6C). This model reflects clinical studies that showed that higher doses of tumor-directed radiation could prolong survival (*4*). These clinical studies, however, were limited to tumors that were more than 1cm away from bowel, which helps to reduce gastrointestinal radiotoxicity, but also limits the types of patients that may benefit from the treatment (*22*). The use of radioprotectors may allow dose-escalated therapy to be available to more patients, which may help improve the outcomes in locally advanced pancreatic cancer. EGLN inhibitors may be an ideal as a first-in-class drug for this purpose, since it exhibits robust radioprotection with little toxicity (*23*), unlike intravenous amifostine, which causes significant hypotension and nausea (*24*).

**Fig 6.**
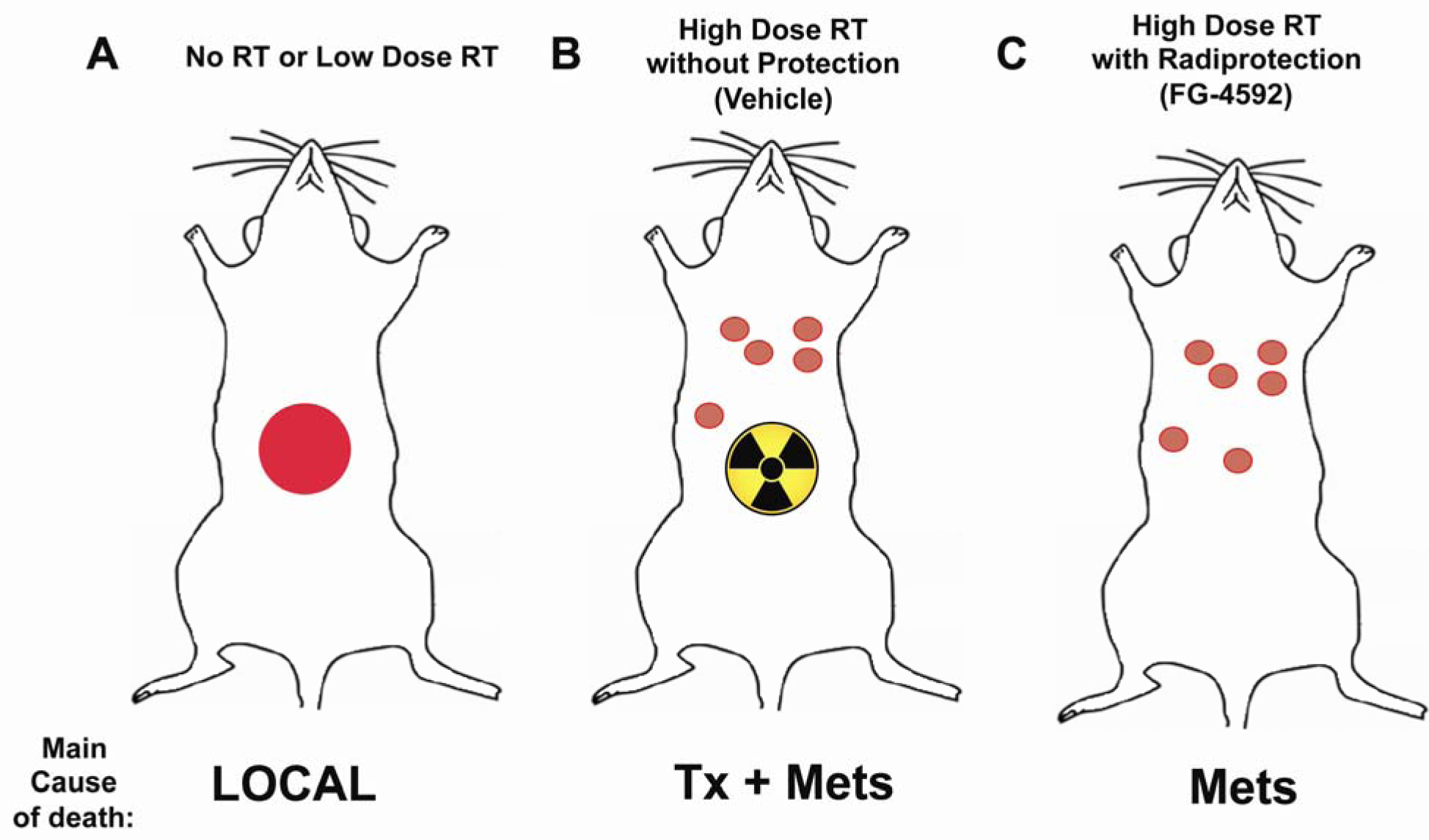
Proposed Pathophysiology of How Radioprotection with RT Improves Survival. (A) *Kras*^LSL/+^;*Trp53*^FL/+^;*Ptf1a*^Cre/+^ (KPC) animals have a locally advanced pancreatic cancer and die primarily from spread of their primary tumor int the abdomen. (B) With high-dose radiation treatments, death due to locally destructive disease is reduced, but is replaced by death from radiation toxicity or metastases. (C) A combination of high-dose radiation and FG-4592 for radioprotection provides local control while preventing toxicity from treatment, which maximizes survival.

KPC animals that received high-dose radiation along with FG-4592 for radioprotection lived longer than their counterparts, but they still eventually succumbed to metastatic burden. This outcome is not surprising, since no chemotherapy was given for microscopic systemic disease. This same shift in causes of treatment failure can be seen in patients. For instance, in the LAP-07 trial, patients that received radiation therapy had improved local control, but exhibited no differences in disease-free survival due to the emergence of metastatic disease that was suboptimally controlled with chemotherapy (*25*). Future efforts preclinical and clinical efforts in LAPC will incorporate modern chemotherapy regimens such as FOLFIRINOX or gemicitabine/abraxane (*26-28*) to further improve survival gains by better controlling metastatic microscopic disease.

We show that it is feasible and desirable to treat pancreatic tumors with fractionated radiation to a definitive dose. Animals that received definitive doses of fractionated radiation (>65Gy) had prolonged survival compared to those that received lower doses (≤65Gy). Moreover, survival between animals that received low dose radiation was indistinguishable from treatments without any radiation at all, which is what has also been reported in clinical trials (*29, 30*). While single fractions of high-dose radiation (such as 10-20Gyx1) are often preferred in preclinical studies due to ease of administration (*31*) these subtherapeutic doses often lead to treatment failure(*16*) and unacceptable toxicity in both preclinical experiments and (*16, 32*) and in clinical trials (*33*). Fractionated (*4, 34*) or stereotactic (*35*) radiation allows sufficient repair of normal tissues (*36*), particularly if paired with a radioprotection, such as FG-4592. Thus, our model should be considered by others to test future radioprotectant interventions with potential for clinical practice.

A common concern with radioprotectors is that they may shield tumors from cytotoxic therapy. Our data clearly demonstrate that EGLN inhibition does not increase intratumoral HIF levels (Fig 2B) nor does it impair treatment outcomes as evidenced by the improved survival in the groups that received FG-4592. We attribute this to the fact that pancreatic tumors are normally quite hypoxic (*37*) and EGLN inhibition cannot raise intratumoral HIF levels any further. In normal intestinal tissue, however, EGLN inhibition can significantly raise HIF expression since there is relatively little hypoxia at baseline (Fig 2A). This differential expression allows for the selective increase of HIF in normal tissues by exploiting the known physiologic properties of the tumor and intestines, respectively. Recent studies demonstrate that HIF might also have tumor suppressive properties in several cancers. For instance, a deletion of *Hif1a* in a model of early pancreatic cancer accelerated the disease, suggesting a tumor suppressor role, while forced expression of Hif2a exhibited tumor suppressive properties in models of lung cancer and sarcoma (*38-41*).

There are some important clinical caveats to our model. We developed KPC animals with a heterozygous deletion of *Trp53* on a C57BL/6J background, which produces a tumor that behaves like locally advanced pancreatic cancer, with morbidity and mortality arising from local progression rather than metastatic burden (Fig 4A and Fig 5A). In clinical practice, only a portion of patients with locally advanced pancreatic cancer have this disease course due to early dissemination (*42*) and inadequate chemotherapy. Newer and more powerful chemotherapy combinations have shown better results in the metastatic disease setting (*26, 28*) and may also find success in the patients with LAPC. Thus, with better systemic treatments, durable local control through dose-escalated radiation may become even more critical in this patient population. Incorporating predictive and prognostic molecular (*3, 43-48*) as well as radiomic biomarkers (*49*) into the clinical translation of these studies will help guide clinical decision making and determine patients most likely ot benefit from dose-escalated radiation therapy. Thus, our finding that intestinal radioprotection by EGLN inhibition enables potentially definitive radiation treatments for LAPC effective should justify clinical trials to test this paradigm for our patients who would otherwise have no options for long-term local control.

## Materials and Methods

### Animals and breeding strategy

Mouse experiments were approved by the UT MD Anderson Institutional Animal Care and Use Committee (IACUC) and were executed in accordance with the NIH guidelines for use and care of live animals. Mice were exposed to 12-hour light/dark cycles and given free access to sterilized powdered food (Prolab Isopro RMH 3000 Irradiated Feed) and sterilized water. C57BL/6J mice were obtained from Jackson Laboratories. *Kras*^LSL/+^ and *Trp53*^FL/FL^;*Ptf1a*^Cre/+^ animals were backcrossed to a pure C57BL/6 background over ten generations and then bred to each other produce the desired *Kras*^LSL/+^;*Trp53*^FL/+^;*Ptf1a*^Cre/+^ animals with 1:4 Mendelian frequency. Genotyping was performed as described previously (*50*).

### Diagnosis, Small Animal Ultrasound and Tumor Measurements

KPC mice with a heterozygous Trp53 deletion typically presented at 16-30 weeks of age. Thus, at 16 weeks of age, mice were subjected to weekly screening for tumors by brief exposure to inhaled anesthesia followed by abdominal palpation. Tumors can palpated from as small as 3mm. Mice with any suspicious lesion on palpation were subjected to ultrasound. Briefly, mice were subjected to 2% isoflurane for anethesia, then shaved with veterinary clippers and briefly treated with epilation cream. Animals were then placed supine onto a warmed ultrasound bed of a Vevo 2100 system (FujiFilm VisualSonics). A 30MHz transducer was used to acquire B-MODE long and short axis acquisitions. In addition a 3D acquisition was also acquired to encompass the entire tumor volume. Length measurements were made on the B-MODE images to assess tumor burden. In addition some 3D volume renderings were done to validate the length measurements. After diagnosis, tumor measurements were made weekly and throughout treatment until death.

### Small Animal Irradiation and Dosimetry

Radiation treatments were administered for 5 consecutive days per week, typically Monday-Friday, followed by a 2 day break (typically Saturday and Sunday) for a total of 15 fractions. Prior to irradiation, mice were anesthetized with 2-3% isoflurane gas mixed with 1L oxygen. Radiation dose was delivered employing X-RAD 225Cx image-guided irradiator from Precision X-Ray, Inc (North Branford, CT). The irradiator combines high-accuracy cone-beam CT (CBCT) imaging (resolution 0.2 mm) and orthovoltage (225 kVp) dose delivery. Each mouse was imaged using the PXi XRAD 225Cx irradiator fitted with a circular 15mm collimator. A cone beam CT prior to irradiation was performed to align the isocenter prior to treatment. The cranial edge of a 15mm circular field was aligned with the curve of the diaphragm on a coronal scout film and then the isocenter was then placed in the mid-abdomen over the tumor, which was defined by ultrasound. Mice received a total of 5Gy per fraction which was treated by equally weighted AP/PA fields (2.5Gy AP and 2.5Gy PA). Imaging dose from cone-Beam CT was negligible (<1 cGy/min).

The dose output of the PXi XRAD 225Cx irradiator was measured with a 0.6cc Farmer chamber FC65G (SN 2247) in the reference geometry (10cm x 10cm) at its isocenter (32.4 cm). The Farmer chamber’s Calibration-Factor (Nk) was provided by the MD Anderson’s Accredited Dosimetry Calibration Laboratory, and is traceable to NIST. Other dosimetry parameters for the Applicators were measured with combination of ion-chamber and film (EBT2, EBT3). The unit’s dose-output constancy is checked prior setting-up animal for irradiation.

### FG-4592 administration

The prolyl hydroxylase inhibitor, FG-4592 (Cayman Chemical, 15294) was administered by oral gavage at a dose of 40mg/kg in 0.5 (w/v)% sterilized methyl cellulose 400 solution (Wako Pure Chemical Industries, Ltd; 133-17815). The initial dose of vehicle or FG-4592 treatment was typically administered 5 days after diagnosis, which was timed to coincide with radiation treatments, if the mice were assigned to receive them. Drug or vehicle dosing continued 4x/week during treatment, which were typically Monday, Wednesday, Friday and Saturday. The treatment was considered complete after after 13 total treatments were administered over 3 weeks. On days when mice received both FG-4592 and radiation, FG-4592 was given 3 hours prior to radiation.

### Immunoblotting

Total protein was extracted by homogenizing tissues in a proprietary tissue extraction buffer (T-PER; ThermoFisher Scientific, Waltham, MA), supplemented with protease and phosphatase inhibitor cocktails (Roche Applied Science, Indianapolis, IN). Lysates were denatured with 4x Laemmli sample buffer (Bio-Rad, Hercules, CA) and fractionated by SDS-PAGE. The proteins were electro-transferred to nitrocellulose membranes using the Trans-blot Turbo system (Bio-Rad, Hercules, CA), then blocked with TBS-T +5% skim milk powder. The blots were probed with primary antibodies diluted in Superblock T20 (ThermoFisher Scientific). HIF-1α (Novus Biologicals, Littleton, CO, NB100-105) was used at 1:500, and HIF-2α/ EPAS1 Novus Biologicals, NB100-122) was used at 1:1000) while β-actin (Cell Signaling Technology, Danvers, MA, #4970) was used at 1:1000. The blots were developed using Clarity^TM^ Western ECL substrate (Bio-Rad). All protein expression data were quantified by Chemi-Doc XRS+ system (BioRad).

### Mouse Histology, Cause of Death, and Necropsy

Mice were subjected to necropsy when they met any euthanasia criteria. Mice were euthanized by CO2 followed by exsanguination via cardiac puncture. Causes of death were assigned after a complete necropsy, but were usually obvious on the initial examination, such intestinal bleeding, outlet obstruction, or diffuse carcinomatosis. For necropsies, tissues were collected and fixed in 10% neutral-buffered formalin. Bones were decalcified in 10% formic acid (Fisher Scientific). Fixed tissues were embedded in paraffin; and tissue sections 4 microns in thickness were placed on slide and stained with hematoxylin and eosin (H&E) and analyzed by a veterinary pathologist (LB).

### Statistical Methods

Descriptive statistics were generated for all mice. Mice who received >65Gy RT were considered as receiving full dose RT and completing treatment. Mice who received <=65Gy RT were considered as having received low dose RT. All mice were included in survival analyses. For mice who underwent necropsy, only mice who received full dose RT were included in comparisons. Sensitivity analyses were performed to demonstrate that exclusion of low-dose RT mice had no impact. Pathologic characteristics were compared using chi square of Fisher’s exact test, where appropriate.

Kaplan-Meier survival curves were generated for all mice, grouped by treatment group. All mice were censored when there were less than 1 mice in all treatment groups (after 50 days). Log-rank test was used to compare overall trends and also for individual stratum comparisons. JMP SAS version 12.0 was used for statistical analyses.

## Acknowledgments

We would like to acknowledge Ronald DePinho and Haoqiang Yang for their generous gift of KPC breeders for our colony. We thank Bob Bast, Anirban Maitra and Joe Herman for their critical review of the data and manuscript.

## Author contributions

T.N.F. and J.M.M. executed all experiments and performed a primary analysis for all mouse experiments with the following exceptions: L.B. performed and analyzed all mouse necropsy experiments, A.D. performed Western blot experiments and L.E.C. performed the biostatistical analysis for all experiments. C.M.T., T.N.F., J.M.M., C.V.K, and G.O.S. designed the radiation experiments and R.C.T and G.O.S. oversaw and performed the dosimetry for the small animal irradiator. The manuscript was written by C.M.T., L.E.C, T.N.F, J.M.M and A.D.

## Supplementary Materials

**Supp Fig 1.**
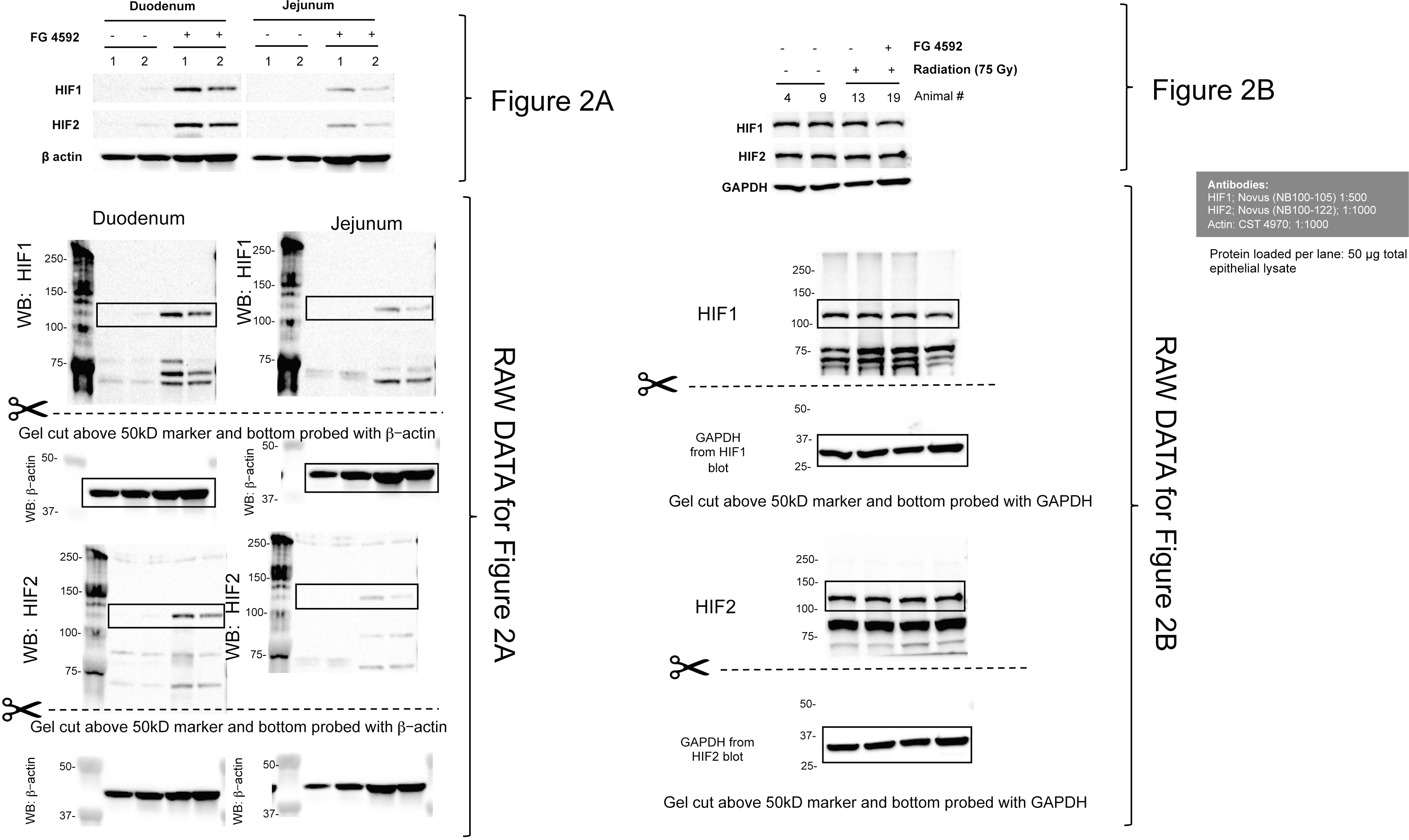
Raw Western Blot Data for Fig 2

## References

1. H. Ying, P. Dey, W. Yao, A. C. Kimmelman, G. F. Draetta, A. Maitra, R. A. DePinho, Genetics and biology of pancreatic ductal adenocarcinoma. Genes Dev 30, 355-385 (2016).

2. L. Rahib, B. D. Smith, R. Aizenberg, A. B. Rosenzweig, J. M. Fleshman, L. M. Matrisian, Projecting cancer incidence and deaths to 2030: the unexpected burden of thyroid, liver, and pancreas cancers in the United States. Cancer Res 74, 2913-2921 (2014).

3. C. A. Iacobuzio-Donahue, B. Fu, S. Yachida, M. Luo, H. Abe, C. M. Henderson, F. Vilardell, Z. Wang, J. W. Keller, P. Banerjee, J. M. Herman, J. L. Cameron, C. J. Yeo, M. K. Halushka, J. R. Eshleman, M. Raben, A. P. Klein, R. H. Hruban, M. Hidalgo, D. Laheru, DPC4 gene status of the primary carcinoma correlates with patterns of failure in patients with pancreatic cancer. J Clin Oncol 27, 1806-1813 (2009).

4. S. Krishnan, A. S. Chadha, Y. Suh, H. C. Chen, A. Rao, P. Das, B. D. Minsky, U. Mahmood, M. E. Delclos, G. O. Sawakuchi, S. Beddar, M. H. Katz, J. B. Fleming, M. M. Javle, G. R. Varadhachary, R. A. Wolff, C. H. Crane, Focal Radiation Therapy Dose Escalation Improves Overall Survival in Locally Advanced Pancreatic Cancer Patients Receiving Induction Chemotherapy and Consolidative Chemoradiation. Int J Radiat Oncol Biol Phys 94, 755-765 (2016).

5. B. D. Kavanagh, C. C. Pan, L. A. Dawson, S. K. Das, X. A. Li, R. K. Ten Haken, M. Miften, Radiation dose-volume effects in the stomach and small bowel. Int J Radiat Oncol Biol Phys 76, S101-107 (2010).

6. S. Stanic, J. S. Mayadev, Tolerance of the small bowel to therapeutic irradiation: a focus on late toxicity in patients receiving para-aortic nodal irradiation for gynecologic malignancies. Int J Gynecol Cancer 23, 592-597 (2013).

7. T. Kamada, H. Tsujii, E. A. Blakely, J. Debus, W. De Neve, M. Durante, O. Jakel, R. Mayer, R. Orecchia, R. Potter, S. Vatnitsky, W. T. Chu, Carbon ion radiotherapy in Japan: an assessment of 20 years of clinical experience. Lancet Oncol 16, e93-e100 (2015).

8. L. Colbert, Moningi, S., Chadha A., Amer, A., Lee, Y., Wollf, R., Vardhachary, G., Fleming, J., Katz, M., Das, P., Krishnan, S., Koay, E., Park, P., Crane, C., Taniguchi, C., Dose escalation with an IMRT technique in 15 to 28 fractions is better tolerated than standard doses of 3DCRT for LAPC. Advances in Radiation Oncology 2, 403-415 (2017).

9. W. G. Kaelin, Jr., P. J. Ratcliffe, Oxygen sensing by metazoans: the central role of the HIF hydroxylase pathway. Mol Cell 30, 393-402 (2008).

10. C. M. Taniguchi, Y. R. Miao, A. N. Diep, C. Wu, E. B. Rankin, T. F. Atwood, L. Xing, A. J. Giaccia, PHD inhibition mitigates and protects against radiation-induced gastrointestinal toxicity via HIF2. Sci Transl Med 6, 236ra264 (2014).

11. M. K. Ayrapetov, C. Xu, Y. Sun, K. Zhu, K. Parmar, A. D. D'Andrea, B. D. Price, Activation of Hif1alpha by the prolylhydroxylase inhibitor dimethyoxalyglycine decreases radiosensitivity. PLoS One 6, e26064 (2011).

12. D. G. Kirsch, P. M. Santiago, E. di Tomaso, J. M. Sullivan, W. S. Hou, T. Dayton, L. B. Jeffords, P. Sodha, K. L. Mercer, R. Cohen, O. Takeuchi, S. J. Korsmeyer, R. T. Bronson, C. F. Kim, K. M. Haigis, R. K. Jain, T. Jacks, p53 controls radiation-induced gastrointestinal syndrome in mice independent of apoptosis. Science 327, 593-596 (2010).

13. W. Qiu, E. B. Carson-Walter, H. Liu, M. Epperly, J. S. Greenberger, G. P. Zambetti, L. Zhang, J. Yu, PUMA regulates intestinal progenitor cell radiosensitivity and gastrointestinal syndrome. Cell Stem Cell 2, 576-583 (2008).

14. P. Vachal, S. Miao, J. M. Pierce, D. Guiadeen, V. J. Colandrea, M. J. Wyvratt, S. P. Salowe, L. M. Sonatore, J. A. Milligan, R. Hajdu, A. Gollapudi, C. A. Keohane, R. B. Lingham, S. M. Mandala, J. A. Demartino, X. Tong, M. Wolff, D. Steinhuebel, G. R. Kieczykowski, F. J. Fleitz, K. Chapman, J. Athanasopoulos, G. Adam, C. D. Akyuz, D. K. Jena, J. W. Lusen, J. Meng, B. D. Stein, L. Xia, E. C. Sherer, J. J. Hale, 1,3,8-Triazaspiro[4.5]decane-2,4-diones as Efficacious Pan-Inhibitors of Hypoxia-Inducible Factor Prolyl Hydroxylase 1-3 (HIF PHD1-3) for the Treatment of Anemia. Journal of medicinal chemistry, (2012).

15. R. Provenzano, A. Besarab, C. H. Sun, S. A. Diamond, J. H. Durham, J. L. Cangiano, J. R. Aiello, J. E. Novak, T. Lee, R. Leong, B. K. Roberts, K. G. Saikali, S. Hemmerich, L. A. Szczech, K. H. Yu, T. B. Neff, Oral Hypoxia-Inducible Factor Prolyl Hydroxylase Inhibitor Roxadustat (FG-4592) for the Treatment of Anemia in Patients with CKD. Clin J Am Soc Nephrol 11, 982-991 (2016).

16. L. Seifert, G. Werba, S. Tiwari, N. N. Giao Ly, S. Nguy, S. Alothman, D. Alqunaibit, A. Avanzi, D. Daley, R. Barilla, D. Tippens, A. Torres-Hernandez, M. Hundeyin, V. R. Mani, C. Hajdu, I. Pellicciotta, P. Oh, K. Du, G. Miller, Radiation Therapy Induces Macrophages to Suppress T-Cell Responses Against Pancreatic Tumors in Mice. Gastroenterology 150, 1659-1672 e1655 (2016).

17. B. W. Fischer-Valuck, L. Henke, O. Green, R. Kashani, S. Acharya, J. D. Bradley, C. G. Robinson, M. Thomas, I. Zoberi, W. Thorstad, H. Gay, J. Huang, M. Roach, V. Rodriguez, L. Santanam, H. Li, H. Li, J. Contreras, T. Mazur, D. Hallahan, J. R. Olsen, P. Parikh, S. Mutic, J. Michalski, Two-and-a-half-year clinical experience with the world’s first magnetic resonance image guided radiation therapy system. Adv Radiat Oncol 2, 485-493 (2017).

18. H. Ying, A. C. Kimmelman, C. A. Lyssiotis, S. Hua, G. C. Chu, E. Fletcher-Sananikone, J. W. Locasale, J. Son, H. Zhang, J. L. Coloff, H. Yan, W. Wang, S. Chen, A. Viale, H. Zheng, J. H. Paik, C. Lim, A. R. Guimaraes, E. S. Martin, J. Chang, A. F. Hezel, S. R. Perry, J. Hu, B. Gan, Y. Xiao, J. M. Asara, R. Weissleder, Y. A. Wang, L. Chin, L. C. Cantley, R. A. DePinho, Oncogenic Kras maintains pancreatic tumors through regulation of anabolic glucose metabolism. Cell 149, 656-670 (2012).

19. J. P. Morton, P. Timpson, S. A. Karim, R. A. Ridgway, D. Athineos, B. Doyle, N. B. Jamieson, K. A. Oien, A. M. Lowy, V. G. Brunton, M. C. Frame, T. R. Evans, O. J. Sansom, Mutant p53 drives metastasis and overcomes growth arrest/senescence in pancreatic cancer. Proc Natl Acad Sci U S A 107, 246-251 (2010).

20. S. R. Hingorani, L. Wang, A. S. Multani, C. Combs, T. B. Deramaudt, R. H. Hruban, A. K. Rustgi, S. Chang, D. A. Tuveson, Trp53R172H and KrasG12D cooperate to promote chromosomal instability and widely metastatic pancreatic ductal adenocarcinoma in mice. Cancer Cell 7, 469-483 (2005).

21. C. K. Haston, Mouse genetic approaches applied to the normal tissue radiation response. Front Oncol 2, 94 (2012).

22. L. E. Colbert, S. Moningi, A. Chadha, A. Amer, Y. Lee, R. A. Wolff, G. Varadhachary, J. Fleming, M. Katz, P. Das, S. Krishnan, E. J. Koay, P. Park, C. H. Crane, C. M. Taniguchi, Dose escalation with an IMRT technique in 15 to 28 fractions is better tolerated than standard doses of 3DCRT for LAPC. Adv Radiat Oncol 2, 403-415 (2017).

23. A. Besarab, R. Provenzano, J. Hertel, R. Zabaneh, S. J. Klaus, T. Lee, R. Leong, S. Hemmerich, K. H. Yu, T. B. Neff, Randomized placebo-controlled dose-ranging and pharmacodynamics study of roxadustat (FG-4592) to treat anemia in nondialysis-dependent chronic kidney disease (NDD-CKD) patients. Nephrol Dial Transplant 30, 1665-1673 (2015).

24. G. A. Hospers, E. A. Eisenhauer, E. G. de Vries, The sulfhydryl containing compounds WR-2721 and glutathione as radio-and chemoprotective agents. A review, indications for use and prospects. British journal of cancer 80, 629-638 (1999).

25. P. Hammel, F. Huguet, J. L. van Laethem, D. Goldstein, B. Glimelius, P. Artru, I. Borbath, O. Bouche, J. Shannon, T. Andre, L. Mineur, B. Chibaudel, F. Bonnetain, C. Louvet, Effect of Chemoradiotherapy vs Chemotherapy on Survival in Patients With Locally Advanced Pancreatic Cancer Controlled After 4 Months of Gemcitabine With or Without Erlotinib: The LAP07 Randomized Clinical Trial. JAMA 315, 1844-1853 (2016).

26. V. Vaccaro, I. Sperduti, M. Milella, FOLFIRINOX versus gemcitabine for metastatic pancreatic cancer. N Engl J Med 365, 768-769; author reply 769 (2011).

27. T. Conroy, F. Desseigne, M. Ychou, O. Bouche, R. Guimbaud, Y. Becouarn, A. Adenis, J. L. Raoul, S. Gourgou-Bourgade, C. de la Fouchardiere, J. Bennouna, J. B. Bachet, F. Khemissa-Akouz, D. Pere-Verge, C. Delbaldo, E. Assenat, B. Chauffert, P. Michel, C. Montoto-Grillot, M. Ducreux, U. Groupe Tumeurs Digestives of, P. Intergroup, FOLFIRINOX versus gemcitabine for metastatic pancreatic cancer. N Engl J Med 364, 1817-1825 (2011).

28. D. D. Von Hoff, T. Ervin, F. P. Arena, E. G. Chiorean, J. Infante, M. Moore, T. Seay, S. A. Tjulandin, W. W. Ma, M. N. Saleh, M. Harris, M. Reni, S. Dowden, D. Laheru, N. Bahary, R. K. Ramanathan, J. Tabernero, M. Hidalgo, D. Goldstein, E. Van Cutsem, X. Wei, J. Iglesias, M. F. Renschler, Increased survival in pancreatic cancer with nab-paclitaxel plus gemcitabine. N Engl J Med 369, 1691-1703 (2013).

29. B. Chauffert, F. Mornex, F. Bonnetain, P. Rougier, C. Mariette, O. Bouche, J. F. Bosset, T. Aparicio, L. Mineur, A. Azzedine, P. Hammel, J. Butel, N. Stremsdoerfer, P. Maingon, L. Bedenne, Phase III trial comparing intensive induction chemoradiotherapy (60 Gy, infusional 5-FU and intermittent cisplatin) followed by maintenance gemcitabine with gemcitabine alone for locally advanced unresectable pancreatic cancer. Definitive results of the 2000-01 FFCD/SFRO study. Ann Oncol 19, 1592-1599 (2008).

30. P. Hammel, F. Huguet, J. L. van Laethem, D. Goldstein, B. Glimelius, P. Artru, I. Borbath, O. Bouche, J. Shannon, T. Andre, L. Mineur, B. Chibaudel, F. Bonnetain, C. Louvet, L. A. P. T. Group, Effect of Chemoradiotherapy vs Chemotherapy on Survival in Patients With Locally Advanced Pancreatic Cancer Controlled After 4 Months of Gemcitabine With or Without Erlotinib: The LAP07 Randomized Clinical Trial. JAMA 315, 1844-1853 (2016).

31. M. Bazalova, G. Nelson, J. M. Noll, E. E. Graves, Modality comparison for small animal radiotherapy: a simulation study. Med Phys 41, 011710 (2014).

32. A. Maeda, Y. Chen, J. Bu, H. Mujcic, B. G. Wouters, R. S. DaCosta, In Vivo Imaging Reveals Significant Tumor Vascular Dysfunction and Increased Tumor Hypoxia-Inducible Factor-1alpha Expression Induced by High Single-Dose Irradiation in a Pancreatic Tumor Model. Int J Radiat Oncol Biol Phys 97, 184-194 (2017).

33. D. Schellenberg, K. A. Goodman, F. Lee, S. Chang, T. Kuo, J. M. Ford, G. A. Fisher, A. Quon, T. S. Desser, J. Norton, R. Greco, G. P. Yang, A. C. Koong, Gemcitabine chemotherapy and single-fraction stereotactic body radiotherapy for locally advanced pancreatic cancer. Int J Radiat Oncol Biol Phys 72, 678-686 (2008).

34. W. A. Hall, L. E. Colbert, D. Nickleach, J. Switchenko, Y. Liu, T. Gillespie, J. Lipscomb, C. Hardy, D. A. Kooby, R. S. Prabhu, J. Kauh, J. C. Landry, The influence of radiation therapy dose escalation on overall survival in unresectable pancreatic adenocarcinoma. J Gastrointest Oncol 5, 77-85 (2014).

35. J. M. Herman, D. T. Chang, K. A. Goodman, A. S. Dholakia, S. P. Raman, A. Hacker-Prietz, C. A. Iacobuzio-Donahue, M. E. Griffith, T. M. Pawlik, J. S. Pai, E. O'Reilly, G. A. Fisher, A. T. Wild, L. M. Rosati, L. Zheng, C. L. Wolfgang, D. A. Laheru, L. A. Columbo, E. A. Sugar, A. C. Koong, Phase 2 multi-institutional trial evaluating gemcitabine and stereotactic body radiotherapy for patients with locally advanced unresectable pancreatic adenocarcinoma. Cancer 121, 1128-1137 (2015).

36. K. A. Mason, L. Milas, N. R. Hunter, M. Elshaikh, L. Buchmiller, K. Kishi, K. Hittelman, K. K. Ang, Maximizing therapeutic gain with gemcitabine and fractionated radiation. Int J Radiat Oncol Biol Phys 44, 1125-1135 (1999).

37. A. C. Koong, V. K. Mehta, Q. T. Le, G. A. Fisher, D. J. Terris, J. M. Brown, A. J. Bastidas, M. Vierra, Pancreatic tumors show high levels of hypoxia. Int J Radiat Oncol Biol Phys 48, 919-922 (2000).

38. M. C. Simon, The Hypoxia Response Pathways - Hats Off! N Engl J Med 375, 1687-1689 (2016).

39. C. Yoon, K. K. Chang, J. H. Lee, W. D. Tap, C. P. Hart, M. C. Simon, S. S. Yoon, Multimodal targeting of tumor vasculature and cancer stem-like cells in sarcomas with VEGF-A inhibition, HIF-1alpha inhibition, and hypoxia-activated chemotherapy. Oncotarget 7, 42844-42858 (2016).

40. M. S. Nakazawa, T. S. Eisinger-Mathason, N. Sadri, J. D. Ochocki, T. P. Gade, R. K. Amin, M. C. Simon, Epigenetic re-expression of HIF-2alpha suppresses soft tissue sarcoma growth. Nat Commun 7, 10539 (2016).

41. J. Mazumdar, M. M. Hickey, D. K. Pant, A. C. Durham, A. Sweet-Cordero, A. Vachani, T. Jacks, L. A. Chodosh, J. L. Kissil, M. C. Simon, B. Keith, HIF-2alpha deletion promotes Kras-driven lung tumor development. Proc Natl Acad Sci U S A 107, 14182-14187 (2010).

42. A. D. Rhim, E. T. Mirek, N. M. Aiello, A. Maitra, J. M. Bailey, F. McAllister, M. Reichert, G. L. Beatty, A. K. Rustgi, R. H. Vonderheide, S. D. Leach, B. Z. Stanger, EMT and dissemination precede pancreatic tumor formation. Cell 148, 349-361 (2012).

43. C. S. Seldon, L. E. Colbert, W. A. Hall, S. B. Fisher, D. S. Yu, J. C. Landry, Chromodomain-helicase-DNA binding protein 5, 7 and pronecrotic mixed lineage kinase domain-like protein serve as potential prognostic biomarkers in patients with resected pancreatic adenocarcinomas. World J Gastrointest Oncol 8, 358-365 (2016).

44. L. E. Colbert, S. B. Fisher, S. Balci, B. Saka, Z. Chen, S. Kim, B. F. El-Rayes, N. V. Adsay, S. K. Maithel, J. C. Landry, W. J. Curran, Jr., High nuclear hypoxia inducible factor 1 alpha expression is a predictor of distant recurrence in patients with resected pancreatic adenocarcinoma. Int J Radiat Oncol Biol Phys 91, 631-639 (2015).

45. L. E. Colbert, A. V. Petrova, S. B. Fisher, B. G. Pantazides, M. Z. Madden, C. W. Hardy, M. D. Warren, Y. Pan, G. P. Nagaraju, E. A. Liu, B. Saka, W. A. Hall, J. W. Shelton, K. Gandhi, R. Pauly, J. Kowalski, D. A. Kooby, B. F. El-Rayes, C. A. Staley, 3rd, N. V. Adsay, W. J. Curran, Jr., J. C. Landry, S. K. Maithel, D. S. Yu, CHD7 expression predicts survival outcomes in patients with resected pancreatic cancer. Cancer Res 74, 2677-2687 (2014).

46. W. A. Hall, A. V. Petrova, L. E. Colbert, C. W. Hardy, S. B. Fisher, B. Saka, J. W. Shelton, M. D. Warren, B. G. Pantazides, K. Gandhi, J. Kowalski, D. A. Kooby, B. F. El-Rayes, C. A. Staley, 3rd, N. Volkan Adsay, W. J. Curran, J. C. Landry, S. K. Maithel, D. S. Yu, Low CHD5 expression activates the DNA damage response and predicts poor outcome in patients undergoing adjuvant therapy for resected pancreatic cancer. Oncogene 33, 5450-5456 (2014).

47. L. E. Colbert, S. B. Fisher, C. W. Hardy, W. A. Hall, B. Saka, J. W. Shelton, A. V. Petrova, M. D. Warren, B. G. Pantazides, K. Gandhi, J. Kowalski, D. A. Kooby, B. F. El-Rayes, C. A. Staley, 3rd, N. V. Adsay, W. J. Curran, Jr., J. C. Landry, S. K. Maithel, D. S. Yu, Pronecrotic mixed lineage kinase domain-like protein expression is a prognostic biomarker in patients with early-stage resected pancreatic adenocarcinoma. Cancer 119, 3148-3155 (2013).

48. S. H. Shin, H. J. Kim, D. W. Hwang, J. H. Lee, K. B. Song, E. Jun, I. K. Shim, S. M. Hong, H. J. Kim, K. M. Park, Y. J. Lee, S. C. Kim, The DPC4/SMAD4 genetic status determines recurrence patterns and treatment outcomes in resected pancreatic ductal adenocarcinoma: A prospective cohort study. Oncotarget 8, 17945-17959 (2017).

49. E. J. Koay, A. M. Amer, F. E. Baio, A. O. Ondari, J. B. Fleming, Toward stratification of patients with pancreatic cancer: Past lessons from traditional approaches and future applications with physical biomarkers. Cancer Lett 381, 237-243 (2016).

50. N. Bardeesy, A. J. Aguirre, G. C. Chu, K. H. Cheng, L. V. Lopez, A. F. Hezel, B. Feng, C. Brennan, R. Weissleder, U. Mahmood, D. Hanahan, M. S. Redston, L. Chin, R. A. Depinho, Both p16(Ink4a) and the p19(Arf)-p53 pathway constrain progression of pancreatic adenocarcinoma in the mouse. Proc Natl Acad Sci U S A 103, 5947-5952 (2006).

